# Multifocal cutaneous neoplastic vascular proliferations in a rainbow boa (*Epicrates cenchria*) collection with boid inclusion body disease

**DOI:** 10.1101/2024.09.12.612643

**Authors:** Anthony Broering Ferreira, Joandes Henrique Fonteque, Jéssica Aline Withoeft, Renata Assis Casagrande, Ubirajara Maciel da Costa, Frank Imkamp, Pauline Göller, Francesca Baggio, Jussi Hepojoki, Udo Hetzel, Anja Kipar

## Abstract

Reports on neoplastic processes in snakes are sparse regardless of their location, origin or behavior. Here, we describe the occurrence of multifocal cutaneous neoplastic processes consistent with hemangioma and hemangioendothelioma, with a differential diagnosis of angiomatosis, in a colony of native Brazilian rainbow boas (*Epicrates cenchria*) which also included animals affected by boid inclusion body disease (BIBD). Thirteen snakes were affected; seven of these had been introduced from other Brazilian sites years earlier, the others had been bred in house but were not offspring of knowingly affected animals. The breeding regime allowed contact between all female and male animals over the years. The cutaneous lesions were first observed over eight years ago, with additional cases detected during the three following years, but no new cases in the last five years. Two affected animals were subjected to a post mortem examination and were found to suffer from peliosis hepatis as one of the additional pathological changes. BIBD was confirmed in five of the eight examined animals, by histology, immunohistology for reptarenavirus nucleoprotein, and multiplex RT-PCR targeting the reptarenavirus S segment. Reptarenavirus infection was also detected in cells in the cutaneous neoplastic processes. PCRs for *Bartonella henselae* and *B. quintana* as well as bacterial DNA in general, performed on a pool of six skin lesions, yielded negative results, ruling out ongoing bacterial infection, like bacillary angiomatosis in humans, of the lesions. The results hint towards an association of reptarenavirus infection and BIBD with neoplastic processes which is worth further investigations.

## Introduction

The literature on the prevalence, types and behavior of neoplastic processes in reptiles is sparse. An older survey of necropsied captive wild animals from the Zoological Society of San Diego collection, USA, came to the conclusion that the prevalence of neoplasms in reptiles (2.19%) was similar to that observed in birds (1.89%) and mammals (2.75%) [1]. A more recent similar study, undertaken at Taipei Zoo in Taiwan, came to a different conclusion and reported a far higher prevalence of neoplastic processes in mammals (8.1%) and birds (4.2%) than in reptiles (1.1%; 5/449) [2]. Two other retrospective studies in the USA focussed on reptiles; for both, the case material comprised necropsied animals and biopsies [3,4]. The first was undertaken in a private exotic species diagnostic service in Washington; it covered case material from a 10-year period (1994-2003) and found a 15% prevalence of neoplastic processes in the snake case material [3]. The second study, from the Philadelphia Zoological Garden (1901-2002), reported an average incidence rate of 3.2% in necropsied snakes, with the highest rate (9.2%) in the period 1992-2002 [4].

The literature also comprises occasional reports on tumors of vascular origin in snakes. Among these are hemangiomas, reported as single masses at various sites, i.e. ovary, cloaca, liver, heart and skin in several snake species of the families *Viperidae* and *Colubridae* [1,3,5–7]. There are also reports on the locally more agressive hemangioendothelioma in snakes, one of these was in the skin of a *Boa constrictor constrictor* [8–10], whereas the aggressive hemangiosarcoma has only been described in the spleen of a corn snake (*Pantherophis guttatus*) and the heart of a Madagascar giant hognose snake (*Leioheterodon madagascariensis*) [11,12]. Another vascular lesion, morphologically similar to hemangioma, is angiomatosis, a condition that is mainly observed in mammals, such as dogs [13] and in humans where it is known to be mainly associated with *Bartonella quintana* or *Bartonella henselae* infection [14].

Neoplastic processes generally develop spontaneously however, they can also be associated with various genetic, hormonal and infectious causes, and senescence [15]. In snakes, evidence of an association of neoplastic processes with other factors is restricted to their apparently sporadic occurrence in animals suffering from boid inclusion body disease (BIBD) [16,17]. BIBD is causally related to reptarenavirus infection [18–20]. Reptarenaviruses (family *Arenaviridae*) are enveloped viruses with a negative-sense single-stranded bisegmented RNA genome with ambisense coding strategy [21,22]: the S segment encodes the glycoprotein precursor and the nucleoprotein (NP), and the L (large) segment encodes the RNA-dependent RNA polymerase and the zinc finger matrix protein (ZP) [22]. Reptarenavirus-positive snakes are frequently co-infected by a combination of genetically distinct segments belonging to different repetarenavirus species, with the amount of L segment species exceeding those of the S segments [23–28]. The transmission of reptarenaviruses can occur both horizontally, through contact or cohabitation, and vertically, to the progeny [23,25,28]. Reptarenavirus infection leads to the formation of characteristic intracytoplasmic inclusion bodies in most cell types of affected snakes [20]; their presence in cells in the peripheral blood or in tissues is considered as the gold standard for the diagnosis of BIBD [26–30].

BIBD is associated with rather unspecific clinical signs such as anorexia, lethargy, and regurgitation, unless in more severe cases, when tremor, opisthotonus, anisocoria, and proprioceptive deficits are observed [31,32]. The disease is also suspected to lead to immunosuppression which would predispose affected animals to fungal and bacterial infections, possibly also to neoplastic processes [16,20]. So far, three cases of neoplastic processes have been reported in boa constrictors with BIBD, two lymphomas and an odontogenic fibromyxoma [33,34]. In two of these, reptarenavirus RNA was detected in the neoplastic tissue, and in the odontogenic fibromyxoma, upon recurrence, the characteristic intracytoplasmatic inclusion bodies were also detected [34].

Here we report the occurrence of multifocal neoplastic vascular skin lesions, in the majority consistent with hemangioma, in a cohort of captive native Brazilian rainbow boas (*Epicrates cenchria*) that also harbored individuals with BIBD.

## Materials and Methods

### Ethics, animals and sample collection

The project was approved by the Animal Use Ethics Committee (CEUA) of CAV-UDESC, No. 4821250722, under SISGEN No. AF52CCD and SISBIO No. 84341-1.

The study was performed on snakes from a private collection in Brazil, housing eight distinct boid species, including three subspecies of *Boa constrictor* and five *Epicrates* species. Animals of different species were housed separately in climate-controlled rooms with exhaust fans where the snakes were kept individually in plastic boxes with coconut chip substrate and plastic hides. Within one species, direct contact between animals was restricted to breeding periods when female animals were housed together with one or two male animals for approximately two months.

The present study was undertaken on a cohort of 29 Brazilian rainbow boas (*Epicrates cenchria*), a species native to the Amazon region of Brazil [35], housed individually in one room. All animals were grossly examined for cutaneous lesions. From 6 snakes, biopsies from 3 cutaneous lesions each were taken, after local anesthetic block with 0.01 mL of 2% lidocaine without vasoconstrictor. The biopsies from one larger lesions of each animal were cut in half; one half each was fixed in 10% buffered formalin for histological examination, the other halves were pooled and frozen at −80 °C in RNA^®^later (Thermo Fisher Scientific). Smaller lesions were fixed in formalin in their entirety.

Two snakes with numerous skin lesions were euthanized upon the owner’s request due to their poor condition, with an overdose of propofol (35 mg/kg) administered into the paravertebral vein. The animals were then subjected to a full diagnostic post mortem examination. Tissue samples were collected from several cutaneous lesions, brain, trachea, lung, heart, esophagus, stomach, intestine, liver, spleen, pancreas, gonads, and kidneys. One set of samples was frozen at −80 °C in RNA^®^later, a second set was fixed in 10% buffered formalin for histological examination.

Information on the individual samples and animals is provided in Supplemental Table S1.

### Histological examination and immunohistology for reptarenavirus nucleoprotein

Tissue specimens for histological examination were trimmed after formalin fixation, processed and routinely embedded in paraffin wax. Sections (3-5 µm) were prepared and stained with hematoxylin and eosin (HE) for histological examination. Additional sections from the livers of the two necropsied animals were routinely stained for the detection of iron (Fe^3+^; Prussian Blue stain). For selected specimens (Supplemental Table S1), consecutive sections were prepared and subjected to immunohistological staining for reptarenavirus nucleoprotein (NP), using the horseradish peroxidase method and a custom-made rabbit polyclonal antibody, as previously described [20,27,30]. Sections incubated with a non-reactive rabbit polyclonal antibody instead of the specific primary antibody served as negative controls.

### Multiplex PCR for reptarenavirus S segments

Tissue specimens frozen in RNA^®^later (Sigma-Aldrich) were thawed and subjected to RNA extraction, RNA quantification, cDNA synthesis and multiplex RT-PCR for the detection of known reptarenavirus S segments, as decribed previously [36]. Since only very weak bands were observed in the gel electrophoresis, two further multiplex RT-PCR approaches were taken. The first was another attempt at the multiplex RT-PCR, with an increased number of cycles, i.e. 42 instead of 38 cycles. In the second, alternative approach, the RT-PCR products from the initial 38 cycle run were purified using the GeneJET PCR Purification Kit (Thermo Fisher Scientific) and subjected to a second round of the multiplex RT-PCR. The PCR products of both approaches were purified using the GeneJET PCR Purification Kit. In addition, after agarose gel electrophoresis, any bands detected at approximately 150 bp, i.e. bands of the amplicon size expected to be specific for reptarenaviruses [36], were excised from the gel and the DNA fragments extracted using the GeneJET Gel Extraction Kit (Thermo Fisher Scientific). Both the RT-PCR purified and the gel extracted amplicons were Sanger sequenced (Microsynth AG, Balgach, Switzerland) using separately the individual primers contained in the multiplex RT-PCR primer mix [36]. BLASTsearch analyses (https://blast.ncbi.nlm.nih.gov/Blast.cgi) served to determine whether the obtained sequences matched with known reptarenavirus S segment sequences.

### Molecular diagnostics for Bartonella infection

DNA was extracted from the pooled skin lesion samples stored in the RNA preserving solution using the QIAsymphony® DSP Virus/Pathogen Kit (Qiagen). A species-specific PCR assay based on a previously published protocol [37], was performed to screen for the presence of *Bartonella henselae* and *B. quintana*. Subsequently, a broad-spectrum bacterial PCR assay was conducted to detect any Bartonella species or other bacterial pathogens present in the samples. This assay amplified a 500 bp fragment of the bacterial 16S rRNA gene using universal primers [38] designed for broad bacterial detection.

## Results

### Animals, clinical signs, frequency and distribution of cutaneous lesions

The *Epicrates cenchria* enclosure housed 29 animals of different origin. Seven individuals had been introduced over some years (but not during the past 10 years) and came from rehabilitation centers. They either were rescued wild animals or had been claimed due to confirmed animal trafficking (i.e. illegal trading of captured wild snakes or illegally bred snakes). The remaining animals had been bred in the facility. All breeding animals had been in contact with each other at least once during the last 15 years, as they were housed together during breeding seasons, when one female was kept with one or two males for 2-3 months, with mating pairs/groups changing for each breeding season.

The age of the animals in the enclosure ranged from 2 to 16 years (7.12 ± 5.7); 17 were male, 12 female. Of the 29 animals in the enclosure, 13 (45%) were found to suffer from multifocal nodular lesions in the skin. The average age of the affected snakes was 14.7 ± 0.95 years; the majority were male (9/13; 69%) four female (31%). Seven of the 13 affected snakes (54%) had been introduced into the colony more than a decade ago; they came from four different rehabilitation centers. All other affected snakes had been bred at the facility; some were siblings or were offspring from the same male; however, they were not bred from the 7 animals introduced from externally. The 16 unaffected snakes, i.e. animals without evidence of cutaneous lesions at the time of sample collection, had been introduced to the enclosure approximately one year earlier. They were young animals (average age: 2.5 ± 0.5 years) entering their first breeding season and were offspring of the affected, older animals.

The owners had first observed cutaneous lesions in individual animals in the enclosure more than eight years ago. The number of affected animals had increased with time over the next 3 years. Since more than five years, no further cases were observed, however, the number of lesions in individual animals increased.

All affected animals exhibited several cutaneous lesions. Overall, the average number of lesions per animal was 16.7 ± 12.4, with a total of 167 lesions counted in the 13 affected animals. Lesions were most frequent in the ventral body region (116/167; 69%), followed by the lateral region (40/167; 24%) and the dorsal region (11/167; 7%). Grossly, they presented as macules (90/167; 54%), endophytic nodules (51/167; 30%) or exophytic nodules (26/167; 16%), ranging from 0.1 cm to 1.2 cm in diameter, with dark red coloration and soft to firm consistency, sometimes with an ulcerated surface and adherent clots and serocellular crusts (Fig 1A-C). Three animals exhibited scars in the ventral region, ranging from 0.3 cm to 0.8 cm in diameter, where nodules had been surgically removed; these had not been subjected to a histological examination.

**Figure 1.**
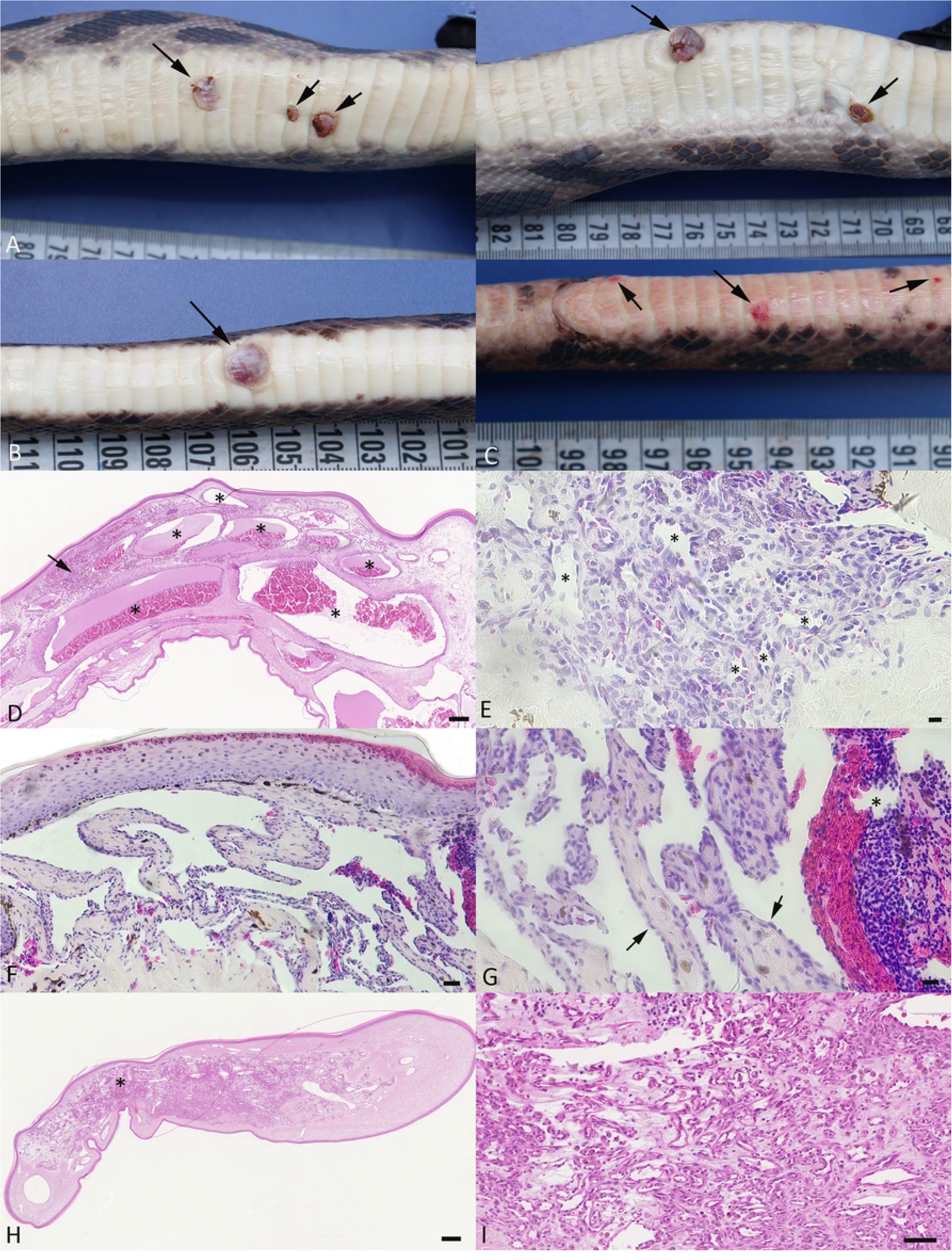
Gross and histological characteristics of the skin lesions. **A-C)** Gross features. **A)** Case 1. Left: Ventral aspect of the body, with 3 variably sized, dark red, slightly elevated nodular lesions. Right: View from lateral, showing exophytic growth of the larger mass (left arrow) and a smaller mass with focal ulceration (right arrow). **B)** Case 3. Ventral aspect of the body, with exophytic, reddish nodular mass (arrow). **C)** Case 4. Ventral aspect of the body, with three slightly elevated, reddish nodular lesions (arrows). **D-I)** Histological features of the skin lesions. **E)** Case 2. Process consistent with cavernous hemangioma. The dermis is expanded by large blood filled cavernae (asterisks) lined by a thin endothelial layer, embedded in an edematous dermal stroma. There are focal leukocyte aggregates (arrow). HE stain. Bar = 125 µm. **E)** Case 6. Process consistent with capillary hemangioma. Aggregate of thin-walled capillaries and small, irregular vascular channels (asterisks), with fibroblasts and a few leukocytes in the surrounding stroma. HE stain. Bar = 20 µm. **F, G)** Case 5. Process consistent with retiform hemangioendothelioma. Dermal lesion comprised of vascular channels lined by differentiated endothelial cells (G: arrows) and supported by fibrous septa. One vascular space contains a fresh thrombus (G: asterisk). HE stains. Bars = 100 µm (E) and 20 µm (F). **H, I)** Case 2. Process suggestive of capillary angiomatosis. **H)** Focal expansion of the dermis by an accumulation of capillaries embedded in fibrous connective tissue. **I)** Closer view of the capillary rich area in H, highlighting the irregular capillaries within an edematous stroma that also contains moderate numbers of leukocytes. HE stains. Bars = 250 µm (G) and 50 µm (H).

In total, an exemplary subset of 24 cutaneous lesions from 8 snakes were subjected to histological examination (Supplementary Table S1). These were represented by irregular structures with an expansion of the subcutis and dermis, covered by a partly folded, often moderately hyperplastic epidermis. The former appeared as non-encapsulated and often poorly delineated accumulations of variably sized, blood-filled vascular structures (Figs. 1D-I, 2). Some represented groups of large vascular clefts and channels embedded in a fibrous to myxoid connective tissue stroma of variable cellularity (Fig. 1D). Vascular structures were lined by a layer of differentiated flat to tombstone-shaped (activated) endothelial cells. The stromal component contained spindle-shaped to polygonal fibroblasts, often with embedded aggregates of siderophages. These histological features were consistent with cavernous hemangioma [13]. Some lesions mainly comprised aggregates of thin-walled capillaries (Figs. 1E, 2A), most consistent with capillary hemangioma [13]. A few were comprised of vascular channels lined by differentiated endothelial cells and supported by fibrous septa (Fig. 1F, G), reminiscent of retiform hemangioendothelioma [13]; these occasional contained thrombi (Fig. 1G). In some masses, irregular accumulations of capillaries embedded in a fibroblast rich, often edematous stroma were observed (Fig. 1H, I); these were reminiscent of capillary angiomatosis [13]. Mitotic figures were generally not detected.

**Figure 2.**
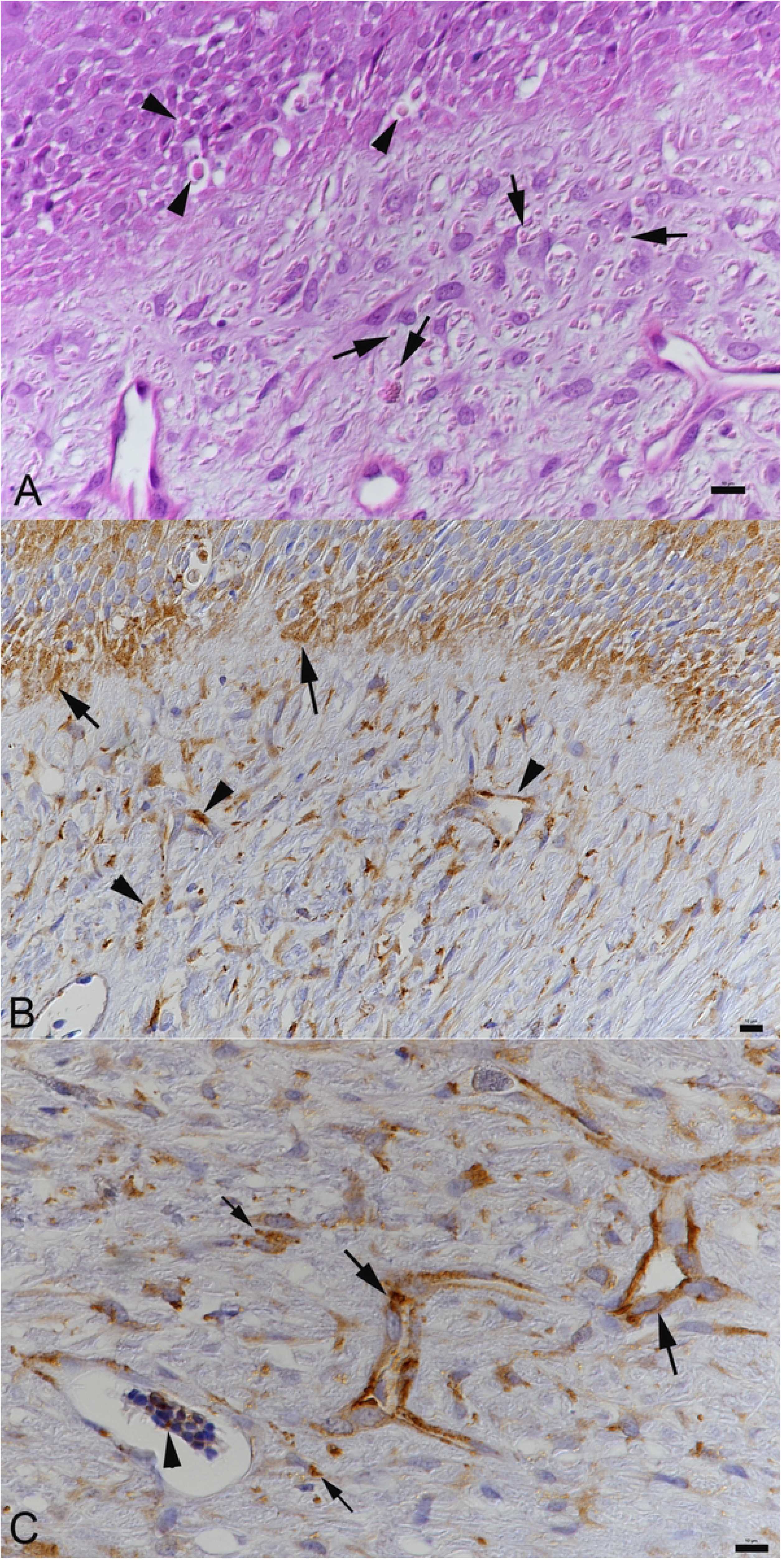
Histological evidence of cytoplasmic inclusion bodies and reptarenavirus nucleoprotein expression in a cutaneous lesion. Case 1**. A**) Close view of the neoplastic process with overlying epidermis. Both epithelial cells in the epidermis (arrowheads) and spindle-shaped cells in the neoplastic process (arrows) exhibit intracytoplasmic inclusion bodies. HE stain. Bar = 10 µm. **B, C**) Immunohistochemistry for reptarenavirus nucleoprotein, hematoxylin counterstain. Bars = 10 µm. **B**) Viral antigen is detected within epithelial cells in the epidermis (arrows) and in cells in the neoplastic process (arrow heads). **C**) Closer view of the neoplastic process. Viral antigen expression is seen in endothelial cells arranged in vascular channels (large arrows) and in spindle shaped cells (small arrows). Arrowhead: positive erythrocyte in an aggregate of blood cells in the lumen of a vascular channel.

In cases 1, 2, 4 and 5, variably sized round cytoplasmic structures reminiscent of the cytoplasmic inclusion bodies were detected in the skin biopsies, both in cells in the dermal neoplastic processes and in the overlying intact epidermis (Fig. 2). They were morphologically identical to the inclusion bodies seen in diverse cell types in snakes with BIBD [20]. Inclusion bodies were not observed in cases 3 and 6 (Supplementary Table S1).

The two necropsied animals, a 16 year old female in moderate body condition, and a 15 year old male in good body condition (cases 7 and 8), had been clinically ill and shown prolonged anorexia, regurgitation, weight loss and apathy, prompting their euthanasia and diagnostic post mortem examination. Both animals exhibited multiple cutanous lesions in particular in the ventral region of the body, but also laterally and, in the case of the male animal (case 8) also dorsally (Supplemental Table S1). The histologically examined skin lesions exhibited morphological features as described above. In both animals, the liver was of diffuse yellowish coloration, with multiple disseminated, partly coalescing dark red subcapsular areas (Fig. 3A). These lesions stretched into the parenchyma and ranged in size from 0.2 to 1.0 cm in diameter. They had a soft consistency and were filled with bloody fluid that oozed out when cut open. Histologically, these lesions presented as poorly delineated areas comprised of blood filled cyst-like structures lined by a thin endothelium, with variable amounts of fibrous connective tissue between and surrounding them and some embedded residual heptic plates (Fig. 3B-E), most consistent with teleangiectasis and/or phlebectatic peliosis hepatis [39,40]. Within and surrounding these areas were abundant siderophages, as confirmed by the Prussian Blue stain for iron (Fe^3+^) (Fig. 3C). The remaining hepatic parenchyma showed multifocal disseminated macrophage aggregates laden with golden pigment (Fig. 3F). This was variably positive for iron (Fig. 3G). Hepatocytes were also found to be laden with iron (Fig. 3C, G), consistent with severe hemosiderosis.

**Figure 3.**
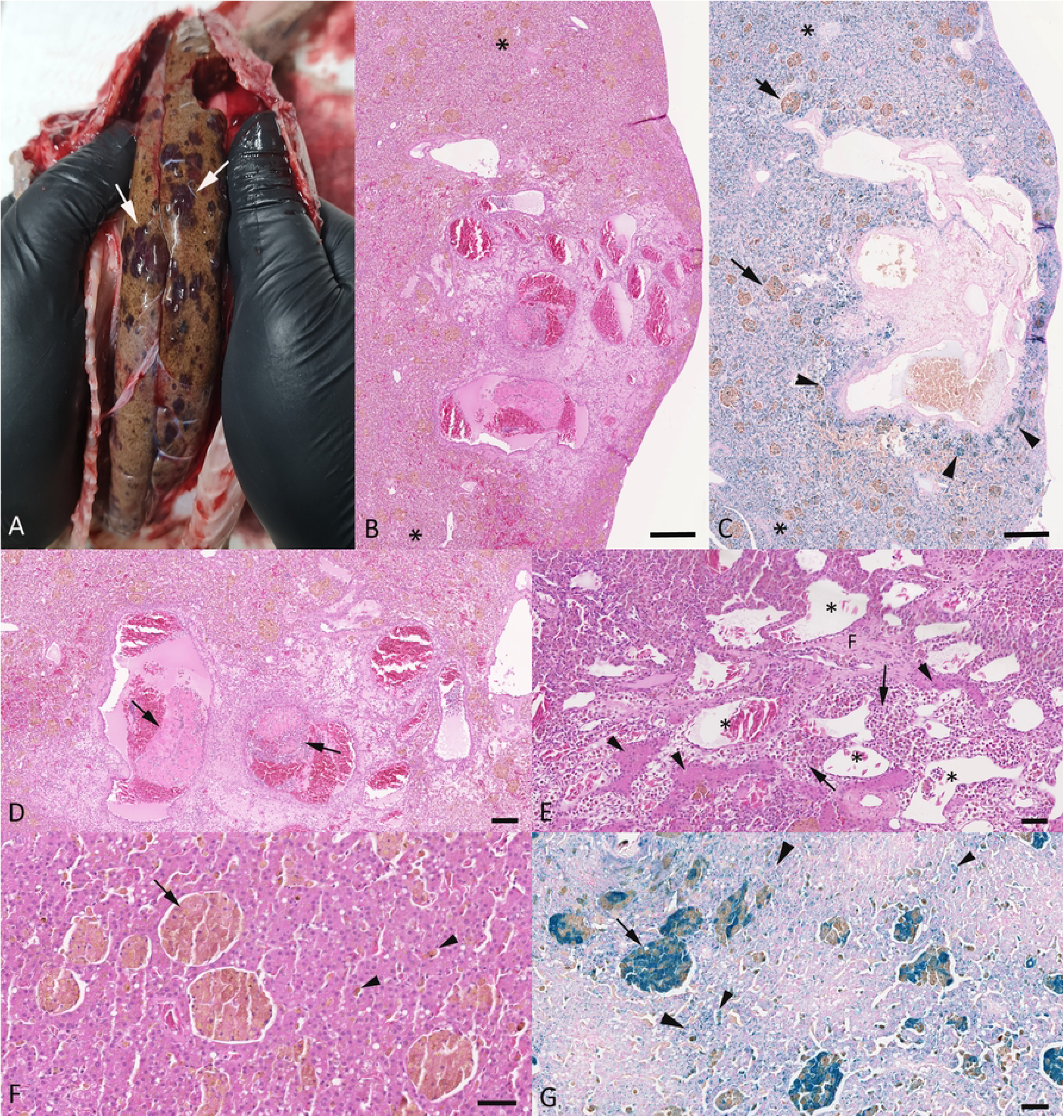
Pathological features in the liver of a female and a male adult *E. cenchria* (cases 7 and 8). **A-D**) Case 7, female animal. **A**) Gross image of the liver *in situ*, with diffuse yellowish coloration. There are multiple disseminated, partly coalescing dark red subcapsular areas (arrows). **B, C**) Focal subcapsular area of teleangiectasis, surrounded by unaffected liver parenchyma (*). The lesion is lined by iron laden macrophages (arrowheads). The latter are also found as variably sized pigment laden aggregates (arrows) in the unaffected parenchyma that also exhibits diffuse cytoplasmic iron deposition in hepatocytes. HE stain (B) and Prussian Blue stain (C). Bars = 250 µm. **D**) Closer view of the teleangiectatic area, with variably sized endothelium lined vascular spaces some of which contain fresh thrombi (arrows). HE. Bar = 125 µm. **E-G**) Case 8, male animal. **E**) Teleangiectatic area with endothelium lined vascular spaces (*) and residual hepatic plates (arrowheads) and islands of fibrous connective tissue (F). Between these structures lie abundant, often pigment laden leukocytes (arrows). HE stain. Bar = 50 µm. **F, G**) Unaffected liver parenchyma. There are multiple, variably sized random aggregates of pigment laden macrophages (F) a lot of which the iron stain confirms to be Fe^3+^ (G). Pigment/iron is also deposited in individual macrophages between hepatic plates (small arrowheads) and in abundant hepatocytes (large arrowheads). HE stain (F) and Prussian Blue stain (G). Bars = 50 µm.

In addition, the female animal (case 7) exhibited a moderate multifocal pyogranulomatous and necrotic hepatitis, consistent with a bacterial etiology, likely due to hematogenous spread of bacteria from the interstine. It also had a small (1 cm in diameter) leiomyoma in the serosa covering the gastric wall.

In the male animal (case 8), the majority of hepatocytes exhibited eosinophilic intracytoplasmic inclusion bodies (Fig. 4A). The latter were also observed in epithelial cells in renal tubules (Fig 4B), testicles, and epididymis (Fig 4C), esophagus, stomach, small and large intestine, adrenal glands and pancreas, and in neurons in the brain (Fig 4D).

**Figure 4.**
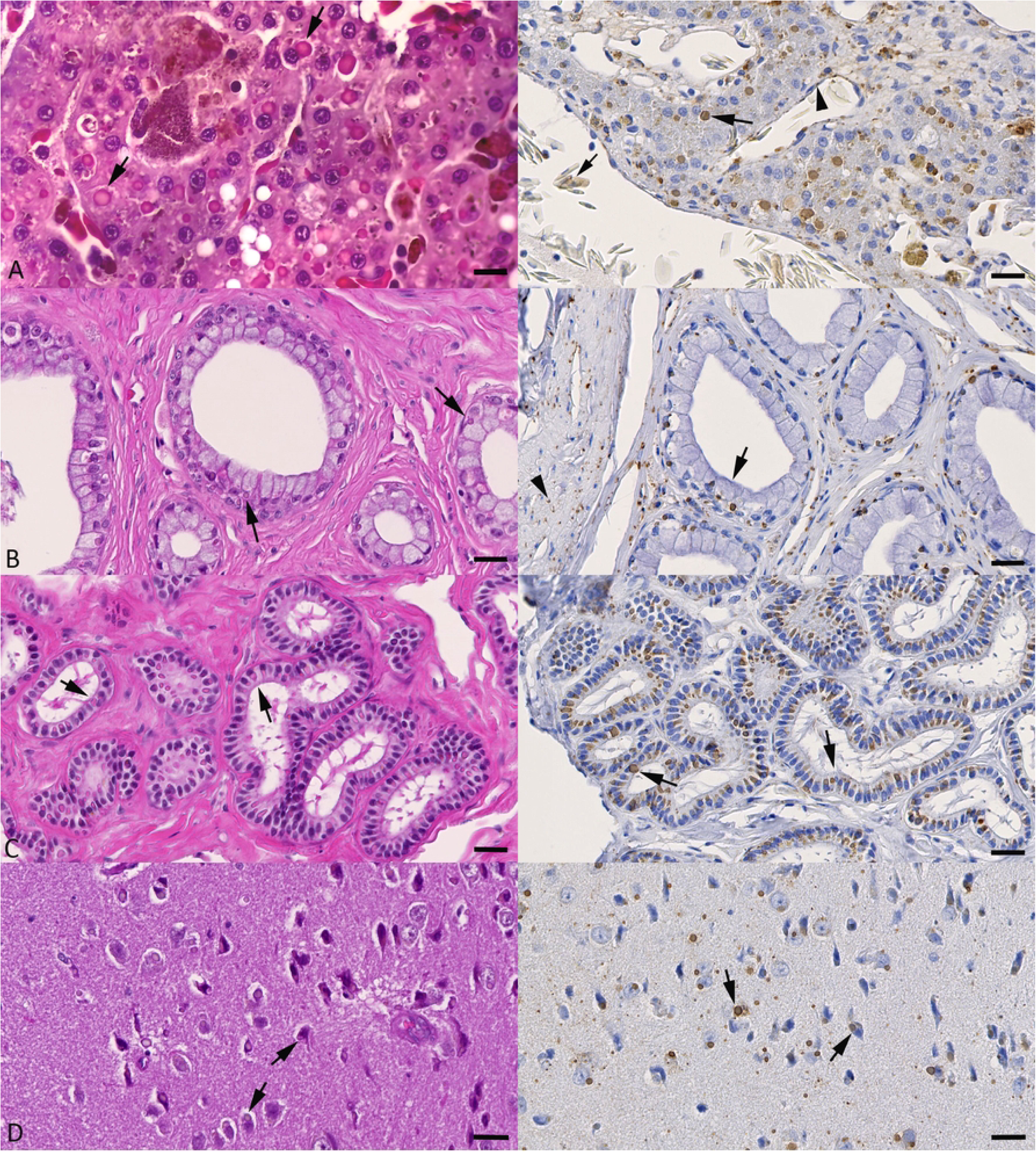
Cytoplasmic inclusion bodies and reptarenavirus nucleoprotein (NP) expression in organs of a male adult *E. cenchria* (case 8). Left column: HE stains; right column: immunohistology for reptarenavirus NP, hematoxylin counterstain. **A**) Liver. Hepatocytes exhibit variably sized, eosinophilic to amphophilic cytoplasmic inclusion bodies (left image: arrows) which are also strongly positive for reptarenavirus NP (right image: arrow). Viral antigen expression is also seen in endothelial cells (arrowhead) and in an erythrocyte (small arrow). **B**) Kidney. Tubular epithelial cells exhibit eosinophilic to amphophilic inclusion bodies (left image: arrows) in the basal cytoplasm. These are strongly positive for viral NP (right image: arrow). Reptarenavirus antigen expression is also seen in interstitial fibrocytes (arrowhead). **C**) Epididymis. The majority of epithelial cells exhibit eosinophilic to amphophilic inclusion bodies (left image: arrows) which are strongly positive for viral NP (right image: arrows). **D**) Brain, cortex. Neurons exhibit variably sized eosinophilic to amphophilic inclusion bodies (left image: arrows). Staining for reptarenavirus NP highlights extensive viral infection in neurons (right image: arrows) and, possibly, glial cells. Bars = 15 µm (A: HE stain) and 20 µm (all other images).

In both animals, all other organs did not exhibit any further histopathological changes.

### Confirmation of reptarenavirus infection and BIBD in a subset of cases

The presence of intracytoplasmic inclusion bodies allowed the diagnosis of BIBD in the male necropsied snake (case 8). BIBD was also indicated by the presence of inclusion bodies in cells in both epidermis and neoplastic processes in the skin biopsies (cases 1, 2, 4 and 5). This prompted an immunohistological examination on tissue sections of some cases, as an attempt to confirm reptarenavirus infection. The results are in line with the histological findings as they highlighted abundant reptarenavirus antigen (nucleoprotein, NP) within endothelial cells and fibroblasts in the vascular lesions as well as in epithelial cells of the overlying epidermis (Fig. 2B, C). We also detected viral antigen in individual circulating blood cells (Fig. 2C), confirming that case 1 was viremic at the time of sampling. Reptarenavirus antigen expression was also shown in cells carrying inclusion bodies in the male necropsied animals (case 8) where it was seen in hepatocytes (Fig. 4A) and endothelial cells; it was also present in esophageal, gastric and intestinal epithelial cells, epithelial cells in renal tubules (Fig. 4B), pancreas, testicle and epididymis (Fig. 4C), cells in spleen and adrenal gland, and in neurons in the brain (Fig. 4D). Erythrocytes were also found to harbor viral antigen (Fig. 4A), confirming that this animal was viremic at the time of death. Immunohistology for viral NP was also performed on sections from the skin biopsy in case 3, and from organs of the female necropsied animal (case 7); this did not yield a positive reaction, hence BIBD was not diagnosed.

Subsequently, we applied our recently established multiplex RT-PCR approach for the detection of reptarenavirus S segments [36] on the samples that had been stored in the RNA preserving solution. The material included a pool of samples from skin biopsies with vascular lesions from six animals (cases 1-6), a liver sample from case 7, and liver, kidney, testicle and skin samples from case 8 (Supplemental Table S1). For all samples, RNA isolation yielded a high RNA concentration (between 80-1800 ng/µL). However, since the samples had accidentally been kept at ambient temperature for one month, we suspected that they had undergone some degree of RNA degradation. Hence, when we obtained very weak bands from the pool of skin biopies and the kidney of case 8 in the agarose gel electrophoresis, we performed a second multiplex RT-PCR run with an increased number of cycles (42 instead of 38 in the original protocol [36] or, alternatively, subjected the purified PCR products from the initial multiplex RT-PCR run to a second RT-PCR run of 38 cycles. The 42-cycle RT-PCR yielded moderately strong bands at the expected length specific for reptarenaviruses, for the pool of skin biopies and the testicle sample of case 8; a very weak band was obtained from the liver sample of case 7. Subsequent Sanger sequencing performed on both purified RT-PCR products and DNA fragments obtained through gel extraction from excised bands, using a subset of the multiplex RT-PCR primers targeting the most common reptarenavirus sequences including the S segments identified in Brazilian boas with BIBD [27] did not yield any sequencing result. When a second RT-PCR run was applied to the PCR products of the initial multiplex RT-PCR, the separation of the RT-PCR products via agarose gel electrophoresis yielded strong bands, corresponding to the expected size for reptarenavirus S segment amplicons, for the pool of skin biopsies and for kidney, testicle and skin samples of case 8. A weak band at the same size was present in the gel for the liver sample of case 8, whereas no band was observed for the liver sample of case 7.

Sanger sequencing of the amplicons from the pool of skin biopsies using the primers included in the multiplex RT-PCR yielded some sequences; the BLAST analysis (https://blast.ncbi.nlm.nih.gov/Blast.cgi) revealed these to be most similar to S5/S5-like(99% identity of an alignment of 101 nucleotides (nt) with primer F5; 100% identity of an alignment of 43 nt with primer R6 [36]) and Tavallinen suomalainen mies virus 2 (TMSV-2, 100% identity of an alignment of 98 nt with primer F4; 99% identity of an alignment of 89 nt with primer R1; 100% identity of an alignment of 49 nt with primer R2 [36]) S segments. Our sequencing attempts on multiplex RT-PCR amplicons obtained from the tissues of case 8 were unsuccessful, even though we confirmed reptarenavirus infection in this histologically BIBD diagnosed animal by immunohistological detection of reptarenavirus NP. In case 7, the negative multiplex RT-PCR result was in agreement with the histological and immunohistological results, i.e. the absence of cytoplasmic inclusion bodies and in particular the lack of reptarenavirus NP expression (Supplemental Table S1).

Molecular diagnostics for the detection of *Bartonella sp*. by PCR, performed on DNA extracted from the pool of skin lesions stored in the RNA preserving solution, did not yield positive results for *B. henselae* and *B. quintana*. The subsequently applied broad-spectrum PCR for bacteria also yielded negative results.

## Discussion

This study reports an outbreak of multifocal benign neoplastic cutaneous lesions in a group of captive Brazilian rainbow boa (*Epicrates cenchria*) in a private collection in Brazil. The skin lesions were widely distributed over the body and grossly and histologically similar, leading to the diagnosis of hemangioma (cavernous and/or capillary) and, in one of the examined lesions, retiform hemangioendothelioma, based on the classification applied to these tumors in dogs and cats [13]. Indeed, this is the first report of cutanous neoplastic vascular processes in this particular snake species, and the second on such a lesion in the skin of snakes [5]. However, some of the lesions carried histological features indicative of angiomatosis; as “capillary angiomatosis”, this is also known as a multinodular entity in the skin of dogs [13].

The presence of multiple cutaneous vascular lesions in all affected snakes is overall unusual. Hemangiomas have only been reported as single processes in snakes so far [1,5,41]; similarly, with the exception of solar-induced dermal hemangiomas [13], cutaneous hemangiomas generally occur as single processes in dogs, as the most frequently affected animal species [13]. Considering the multiplicity of the lesions and the observation that all cases occurred within a limited time frame, i.e. 3 years, without new cases in the subsequent 5 years, the thought arises that the condition is transmissible. This is where the differential diagnosis of “angiomatosis” comes back into play. Humans can suffer from a skin condition called “bacillary angiomatosis”, an angioproliferative lesion induced by *Bartonella henselae* and/or *B. quintana* infection [14]. *Bartonella* sp. are Gram-negative bacteria that primarily infect erythrocytes; they have animal reservoirs, are globally distributed and can be considered as virtually ubiquitous in mammals [42]. However, they have also been detected in reptiles, i.e. sea turles (*Caretta caretta*) [43]. The reservoir of *B. henselae* are cats, although the bacterium has also been identified in the blood of harbor porpoises (*Phocoena phocoena*) and sea turles [14,43,44], while humans are the reservoir of *B. quintana* [14]. Infection of human patients occurs after trauma, such as scratches and bites from cats for *B. henselae*, or through vectors like mites, lice, or fleas [42,45]. Interestingly, infection with *B. henselae* can also be associated with peliosis hepatis in affected people [14], as the bacteria align along and infect endothelial cells in vascularised tissues and induce their proliferation [42]. Lesions due to both *B. henselae* and *B. quintana* are primarily seen in immunocompromised patients [14]. Considering that other *Bartonella sp.* have, for example, rodents as reservoirs [14], one could suspect *Bartonella* infection as the potential cause of the skin lesions in the snakes in the present study. After all, not only do snakes feed on rodents, they are also frequently infested with mites [46] which are also suspected as a source of transmission of reptarenaviruses [16], the causative agents of BIBD. Indeed, we found 5 of the 8 examined animals to be suffering from BIBD, a disease that is believed to be associated with immunosuppression or at least an altered immune status [16,47]. Maintenance of the lesions in humans depends on the presence of bacteria [42]. We did not detect any evidence of bacterial colonisation in any of the skin and peliosis hepatis samples, and the PCRs for *B. henselae* and *B. quintana* were both negative; even a PCR for bacteria in general yielded a negative result. We therefore have no clear evidence that *Bartonella sp*. or bacteria in general are the cause of the lesions. On the other hand, in principle we cannot exclude that we are dealing with lesions that have persisted after the animals have overcome the causative bacterial infection. Furthermore, we also have to consider the option that the DNA quality in the samples was too poor, due to accidental suboptimal storing. In any case, this is an area that warrants further investigation.

In the studied snake cohort, while often widespread on the body, the cutaneous lesions were most numerous in the ventral region, while in snakes cutaneous neoplastic processes appear to generally occur most frequently in the dorsolateral region [48]. Coincidentally, the ventral abdomen is the main site of canine solar-induced dermal hemangioma [13]; such an association can however be excluded in the snakes as they are housed indoors without exposure to sunlight, the environmental temperature in the facility is maintained by thermal plates that are not in contact with the animals (personal communication). Assuming an infectious nature of the skin lesions, they might also have taken advantage of the fact that over the years, all male breeding animals had been housed with their female counterparts, and frequently as two males with one female.

The present study is also the first to describe BIBD and reptarenavirus infection in *Epicrates cenchria* in a collection in their native country, as so far the only cases of BIBD in *E. cenchria* came from collections in the USA [29,49]. The genus *Epicrates* belongs to the family Boidae and shares a monophyletic clade with three other genera present in Brazil: *Boa, Corallus* and *Eunectes* [35,50]. It is now the second boid species native to Brazil in which BIBD is confirmed, as we previously reported the disease in four *Boa constrictor constrictor* snakes held in captivity in Porto Alegre [27]. We diagnosed BIBD using complementary approaches. In 4 cases (cases 1, 2, 4, 5) of which only skin lesions were available, the histological examination detected the pathognomonic intracytoplasmic inclusion bodies in cells in the vascular neoplastic lesions and the overlying epidermis. Immunohistology for reptarenavirus NP, performed on sections of one skin lesion (case 1), confirmed infection of epidermal epithelial cells as well as endothelial cells and fibroblasts within the neoplastic process. Our recently established multiplex RT-PCR, designed for the detection of most known reptarenavirus S segment species [36] served in attempts to provide additional molecular proof of reptarenavirus infection. Specifically, after two RT-PCR runs on the pooled skin lesion samples, we identified through Sanger sequencing S species (nearly) identical to S5/S5-like and TMSV-2 reptarenaviruses S segments. The S5 segment was formerly identified in *Boa constrictors* from Lousiana, USA [23], and the S5-like and TMSV-2 segments were detected in captive boa constrictors from European breeding collections, specifically in Germany and Switzerland [25,26,51]. This could in principle suggest that these reptarenavirus species are widespread and might have been introduced to captive snake collections in Europe and the USA by traded wild-caught snakes carrying the infections. However, given the high similarity with reptarenavirus sequences identified only in European captive snakes [25,26,51] and not in snakes from Brazil [27], one needs to consider laboratory cross-contamination, in particular since amplicons referable to reptarenavirus S segments sequences were only detected after two subsequent RT-PCR runs, increasing the risk of such an event. In any case, using immunohistology for viral NP we could provide definite proof of reptarenavirus infection also in one of the two necropsied snakes (case 8) in which we failed to detect reparenaviruses S segment sequences. Therefore, further studies including a next generation sequencing (NGS) approach on higher quality material from this snake cohort would be required to identify the reptarenaviruses present in the collection.

In a recent NGS-based study, conducted on four Brazilian captive indigenous snakes with BIBD, we identified two novel reptarenavirus S segment species, i.e. Porto Alegre virus (PAV) and Aramboia virus (ArBV) [27]. Our multiplex RT-PCR was designed to also detect these novel species [36] but the test did not pick up these reptarenaviruses in the *E. cenchria* samples analyzed. Curiosly, the metatranscriptomic studies done on snakes with BIBD so far have resulted in the discovery of approximately 30 reptarenavirus L segments while less than 15 S segments have thus far been identified [23,27]. Due to the bisegmented nature of reptarenaviruses and the genetic divergence between the identified L segments, it appears likely that we only know the S segment sequence for <50% of reptarenaviruses. Based on this rationale, it is possible that our RT-PCR approach could have failed in confirming reptarenavirus infection in the studied animals, suggesting that the repertoire of reptarenavirus segments circulating in the natural habitats of boas would be broader than that observed in captive collections. Subjecting more suitable material from animals of the examined cohort and of other native snake cohorts to metatranscriptomic analyses might therefore represent an appropriate systematic approach to challenge this hypothesis and gain further insight into the geographical differences in snake reptarenaviromes.

Interestingly, in line with the above, the formerly reported cases of BIBD in *Epicrates cenchria* were diagnosed solely on the basis of the presence of inclusion bodies in HE stained sections [29,49], as they resulted reptarenavirus negative when a previously assessed method for reptarenavirus detection by RT-PCR was employed [23,29,49]. In the same work, the authors could not detect reptarenavirus NP by immunohistology or immunoblotting, leading to their hypothesis that the inclusion bodies in *E. cenchria* might be caused by unknown reptarenavirus variant(s), or, alternatively, from a protein unrelated to BIBD [29].

We have no means to identify the potential source of the reptarenavirus infection in the cohort. However, introduction from outside Brazil can be excluded (personal communication). Also, among the BIBD positive snakes was at least one that had been introduced from outside (and possibly represented a wild caught animal). Considering the fact that we have previously confirmed BIBD in wild snakes, albeit from Costa Rica [30], and had a similar scenario in boa constrictors with BIBD from Brazil [27], it cannot be excluded that the source was an infected wild *Epicrates cencheri*. All other animals could then have been infected during breeding, or vertically.

So far, three cases of neoplastic processes in snakes with BIBD have been reported, all in *Boa constrictor*: a precardial lymphoma with leukemia, an intestinal lymphoma, and an odontogenic fibromyxoma [33,34]. The histological examination did not reveal viral inclusion bodies in either neoplastic process, however, an RT-PCR for arenavirus performed at a commercial provider yielded a positive result in two cases [34]. Two years later, the odontogenic fibromyxoma had recurred and was examined histologically. At this point, inclusion bodies were detected in lymphocytes, hepatocytes and neoplastic cells, and arenavirus infection was again confirmed by RT-PCR [34]. This and the present cases confirm reptarenavirus proliferation in neoplastic processes. Given the very broad target cell spectrum of the viruses, including both labile (such as hematopoietic cells [47]) and stable and irreversibly postmitotic cells (such as hepatocytes and neurons, respectively), this is not surprising and could be considered as coincidental. There is so far no hard evidence that reptarenaviruses have oncogenic potential. However, arenaviruses, as practically all viruses, need to suppress the immune system to facilitate their replication, and thus it could be speculated that the resulting immunosuppression might contribute to tumorigenesis during persistent infection.

The two necropsied animals had been clinically ill, with rather non-specific signs (prolonged anorexia, regurgitation, weight loss and apathy). In the male animal (case 8), these could in principle be attributed to BIBD [16]. The female animal (case 7), without BIBD, suffered from a pyogranulomatous and necrotic hepatitis, which is indicative of a septic process; this could also explain the clinical signs. In addition, both animals also exhibited severe hepatic hemosiderosis, an alteration so far only reported rarely in snakes, in random cases, a corn snake with hepatotoxic changes due to the antiparasitic agent halofuginone [52], a kingsnake months after the surgical removal of an undifferentiated sarcoma [53], and a *Boa constrictor constrictor* with a glial or ependymal neoplasm [54]. They also showed teleangiectasis/peliosis hepatis which would be consistent with prior liver damage [39,40].

To some extent the results of the present study raise more questions than answers, as it describes multifocal neoplastic processes of vascular origin and potentially infectious nature in snakes with and without BIBD and confirmed reptarenavirus infection. Similar multifocal neoplastic conditions have so far not been reported in snakes but is morphologically reminiscent of bacillary angiomatosis in humans; it might not be possible to determine whether there is indeed a bacterial etiolgy involved, as no new cases have been observed for a longer time period. On the other hand, one could consider a causative role of reptarenaviruses in this scenario. Persistent reptarenavirus infection could dampen the immune system and, if not allowing bacterial infection, promote neoplastic transformation of infected cells, most likely endotheial cells in the present cases. The failure to detect reptarenaviruses in some cases could then indicate that the animals have overcome the infection at some point after onset of the neoplastic processes. Further studies would be needed to challenge or confirm these hypotheses.

## Conclusion

This study documents, for the first time, cutaneous neoplastic vascular processes in *Epicrates cenchria*, represented by multifocal hemangiomas and hemangioendothelioma in thirteen animals. The multiplicity and distribution of the lesions and the time-limited occurrence of cases suggest a complex pathogenesis, possibly involving immunosuppression and persistent infection by reptarenaviruses identified in the studied population, which may predispose the animals to tumorigenesis. However, further studies are necessary to understand the pathogenesis of this neoplastic condition as well as the speculatated tumorigenic role of reptarenaviruses.

## Acknowledgements

The authors are grateful to the technical team in the histology and molecular biology laboratories at the contributing institutions, for excellent technical support.

